# ExposoGraph: An Interactive Platform for Carcinogen Bioactivation and Detoxification Pathway Visualization

**DOI:** 10.64898/2026.03.22.713456

**Authors:** Julhash U. Kazi, Kenneth J. Pienta

## Abstract

**Background:** Despite extensive cataloging of carcinogenic exposures by the International Agency for Research on Cancer (IARC) and pharmacogenomic variation by resources such as PharmVar and CPIC, few platforms unify exposure, metabolic activation and detoxification, DNA damage, and genetic annotation within a single interactive visualization framework. This gap limits systematic evaluation of gene-environment interactions in cancer risk assessment.

**Methods:** We developed the Carcino-Genomic Knowledge Graph, ExposoGraph, an interactive knowledge-graph platform for carcinogen metabolism and DNA damage pathways. The reference graph integrates curated data and annotations from IARC, KEGG, PharmVar, CPIC, CTD, and supporting literature/resources. The current reference graph contains 96 nodes across 5 entity types (Carcinogens, Enzymes, Metabolites, DNA Adducts, and Pathways) and 102 edges across 6 relationship types (activates, detoxifies, transports, forms adduct, repairs, and pathway).

**Results:** The first-generation reference graph captures metabolic activation and detoxification pathways for 9 carcinogen classes spanning 15 index carcinogens. It represents 36 enzymes across Phase I activation (n=14), Phase II conjugation and detoxification (n=14), Phase III transport (n=3), and DNA repair (n=5). Interactive exploration supports carcinogen-class filtering, node- and edge-type filtering, metadata-based search, and detailed hover/detail views with provenance and pharmacogenomic annotations. The androgen branch highlights cross-pathway connectivity by linking androgen metabolism to estrogen quinone formation and DNA adduct generation through CYP19A1-mediated aromatization and downstream catechol estrogen chemistry. In the optional androgen-focused extension, additional receptor, tissue, and variant context further connects this branch to androgen receptor signaling and genotype-specific annotations.

**Conclusions:** ExposoGraph provides a first-generation integrated, interactive framework linking carcinogenic exposures to metabolic fates and genetic modulators. The platform supports hypothesis generation for gene-environment interaction studies and may inform future individualized risk modeling, while remaining a research-use framework rather than a clinically validated risk-assessment tool.

## INTRODUCTION

Chemical carcinogenesis represents a major and modifiable component of the global cancer burden. The International Agency for Research on Cancer (IARC) has classified over 120 agents as Group 1 human carcinogens through its Monographs programme, which systematically evaluates the carcinogenic hazard of chemical, physical, and biological agents (1). Yet the impact of these exposures on any individual is profoundly shaped by genetic variation in the enzymes that metabolize them. Wild’s seminal exposome concept recognized that the totality of environmental exposures, from conception onward, collectively contributes to disease risk in ways that cannot be captured by genomic data alone (2). Recent analyses suggest that environmental factors explain up to 17% of mortality variation, far exceeding the less than 2% attributable to polygenic risk scores (3). This asymmetry underscores the continued need for tools that integrate exposure biology with individual genetic architecture.

Xenobiotics are foreign chemical substances found within an organism or ecosystem that are not naturally produced or expected to be present, such as drugs, environmental pollutants, pesticides, and industrial chemicals (4). The majority of chemical carcinogens are pro-carcinogens that require metabolic activation by xenobiotic-metabolizing enzymes to form ultimate carcinogens capable of covalent DNA modification. This biotransformation involves a coordinated series of enzymatic steps: Phase I activation, predominantly mediated by cytochrome P450 (CYP) enzymes that introduce reactive functional groups; Phase II conjugation by glutathione S-transferases (GSTs), N-acetyltransferases (NATs), UDP-glucuronosyltransferases (UGTs), sulfotransferases (SULTs), and catechol-O-methyltransferase (COMT) that typically render metabolites more hydrophilic and excretable; Phase III transport by ATP-binding cassette (ABC) transporters that export conjugated metabolites from the cell; and DNA repair mechanisms that correct adducts before they become fixed mutations. The dynamic balance between activation and detoxification pathways ultimately determines an individual’s net carcinogenic risk from a given exposure (5).

Pharmacogenomic research over the past three decades has extensively characterized functional genetic variation in carcinogen-metabolizing genes (6, 7). The Pharmacogene Variation (PharmVar) Consortium catalogs over 660 star alleles across more than 30 pharmacogenes, providing a standardized nomenclature for clinically relevant variants (8). The Clinical Pharmacogenetics Implementation Consortium (CPIC) publishes peer-reviewed guidelines for translating diplotypes into functional phenotypes and, where applicable, therapeutic recommendations (9-11). The epidemiological consequences of this variation are well established. For example, the GSTM1-null deletion, present in approximately 50% of Caucasian populations, is associated with increased lung cancer risk (OR = 1.46, 95% CI: 1.07-2.00) (12). NAT2 slow acetylation confers an elevated risk of bladder cancer from aromatic amine exposure (OR = 1.4, 95% CI: 1.2-1.6) (13). The CYP1A1 Ile462Val polymorphism modulates susceptibility to polycyclic aromatic hydrocarbon (PAH)-related malignancies (OR = 1.22) (14). Critically, these risks are multiplicative: individuals carrying both CYP1A1*2C (high activity) and GSTM1-null genotypes exhibit an OR of 4.67 (95% CI: 2.00-10.9) for lung cancer in non-smokers, illustrating the power of gene-gene-environment interactions (15).

Several existing tools address individual components of this landscape. PharmCAT extracts pharmacogenomic variants from variant call format (VCF) files and generates CPIC-based clinical reports for drug-gene pairs (16, 17). The Comparative Toxicogenomics Database (CTD) curates over 45 million chemical-gene-disease relationships from the published literature (18). The Kyoto Encyclopedia of Genes and Genomes (KEGG) provides reference metabolic pathway maps, including those for xenobiotic metabolism and chemical carcinogenesis (19). Cytoscape offers a powerful, general-purpose network visualization environment for biomolecular interaction data (20). However, none of these tools integrates carcinogen metabolic activation pathways with pharmacogenomic variation in an interactive, web-accessible format specifically designed for cancer risk assessment. Researchers seeking to understand how an individual’s genetic profile modulates their risk from a specific carcinogen exposure must manually synthesize information across multiple databases—a time-consuming process that impedes both hypothesis generation and clinical translation.

Here we present the ExposoGraph, an interactive Carcino-Genomic Knowledge Graph, D3 HTML-based force-directed network visualization that integrates carcinogen metabolism, genetic variation, and DNA damage pathways into a unified platform. The ExposoGraph maps the metabolic journey for 9 classes of chemical carcinogens through 36 metabolizing enzymes to their ultimate DNA adduct products, enabling researchers and clinicians to visualize how individual genetic profiles modulate cancer risk from specific environmental exposures. The platform is freely accessible from a web browser from web browser (https://exposograph.streamlit.app) or can be used locally by following the instructions (https://exposograph.readthedocs.io).

## METHODS

### Data sources and curation

The ExposoGraph was constructed through systematic curation of data from seven primary sources. Carcinogen classifications were curated from a subset of the IARC Monographs programme, which provides the most authoritative evaluation of carcinogenic hazard for chemical, physical, and biological agents (1). Metabolic route information was cross-referenced with the KEGG Pathway Database, specifically the xenobiotic metabolism by cytochrome P450 pathway (hsa00980), steroid hormone biosynthesis (hsa00140), chemical carcinogenesis—DNA adducts (hsa05204), and chemical carcinogenesis—reactive oxygen species (hsa05208) reference pathways (19). Allele nomenclature and functional classifications for supported pharmacogenes were obtained from the PharmVar Consortium database (8). Genotype-to-phenotype translation rules and activity score methodology followed CPIC standards for guideline-backed genes (9-11). Chemical-gene interaction data were supplemented from the Comparative Toxicogenomics Database (CTD) (18). Tissue-specific expression context for metabolizing enzymes was derived from GTEx Portal gene-expression references (21). Enzyme-specific functional data, including variant-function annotations and supporting evidence, were curated from primary literature indexed in PubMed.

### Knowledge graph architecture

The ExposoGraph reference graph comprises five node types and six edge types, organized to represent metabolic paths from carcinogen exposure to DNA damage (Figure 1). The five node types are: Carcinogen (n = 15), representing curated chemical carcinogens and endogenous hormone-related carcinogen contexts; Enzyme (n = 36), representing xenobiotic-metabolizing enzymes and DNA repair proteins; Metabolite (n = 28), representing intermediate and ultimate reactive metabolites; DNA_Adduct (n = 11), representing covalent DNA modifications produced by the ultimate carcinogens; and Pathway (n = 6), representing KEGG reference pathways and curated pathway-context nodes that provide broader metabolic context. The total graph contains 96 nodes. Six directed edge types encode the biological relationships between nodes. ACTIVATES edges connect enzymes to the metabolites they produce from parent carcinogens through Phase I oxidation, reduction, or hydrolysis. DETOXIFIES edges connect Phase II enzymes to the metabolites they render less reactive through glutathione conjugation, glucuronidation, acetylation, sulfation, or related detoxification steps. TRANSPORTS edges connect ABC transporters to conjugated metabolites they export from the cell. FORMS_ADDUCT edges connect reactive metabolites to the specific DNA adducts they produce. REPAIRS edges connect DNA repair enzymes to the adducts they remove or correct. PATHWAY edges map entities to their corresponding pathway nodes. The total graph contains 102 edges. Carcinogens are organized into nine classes reflecting their chemical structure and metabolic behavior: polycyclic aromatic hydrocarbons (PAH; benzo[a]pyrene, DMBA), heterocyclic amines (HCA; PhIP, MeIQx), aromatic amines (4-aminobiphenyl, benzidine), nitrosamines (NNK, NDMA), mycotoxins (aflatoxin B1), estrogens (17-beta-estradiol), androgens (testosterone, 5-alpha-dihydrotestosterone), solvents (benzene, vinyl chloride), and alkylating agents (ethylene oxide). Enzymes are categorized by functional phase: Phase I metabolism (n = 14, including CYP1A1, CYP1B1, CYP1A2, CYP2A6, CYP2A13, CYP2E1, CYP3A4, CYP17A1, SRD5A1, SRD5A2, CYP19A1, CYP3A5, AKR1C3, and HSD3B2), Phase II metabolism (n = 14, including EPHX1, GSTM1, GSTT1, GSTP1, NAT1, NAT2, SULT1A1, UGT1A1, UGT2B7, UGT2B17, UGT2B15, NQO1, COMT, and AKR1C2), Phase III transport (n = 3; ABCB1, ABCG2, ABCC2), and DNA repair (n = 5; OGG1, XRCC1, ERCC2/XPD, XPC, and MGMT).

**Figure 1.**
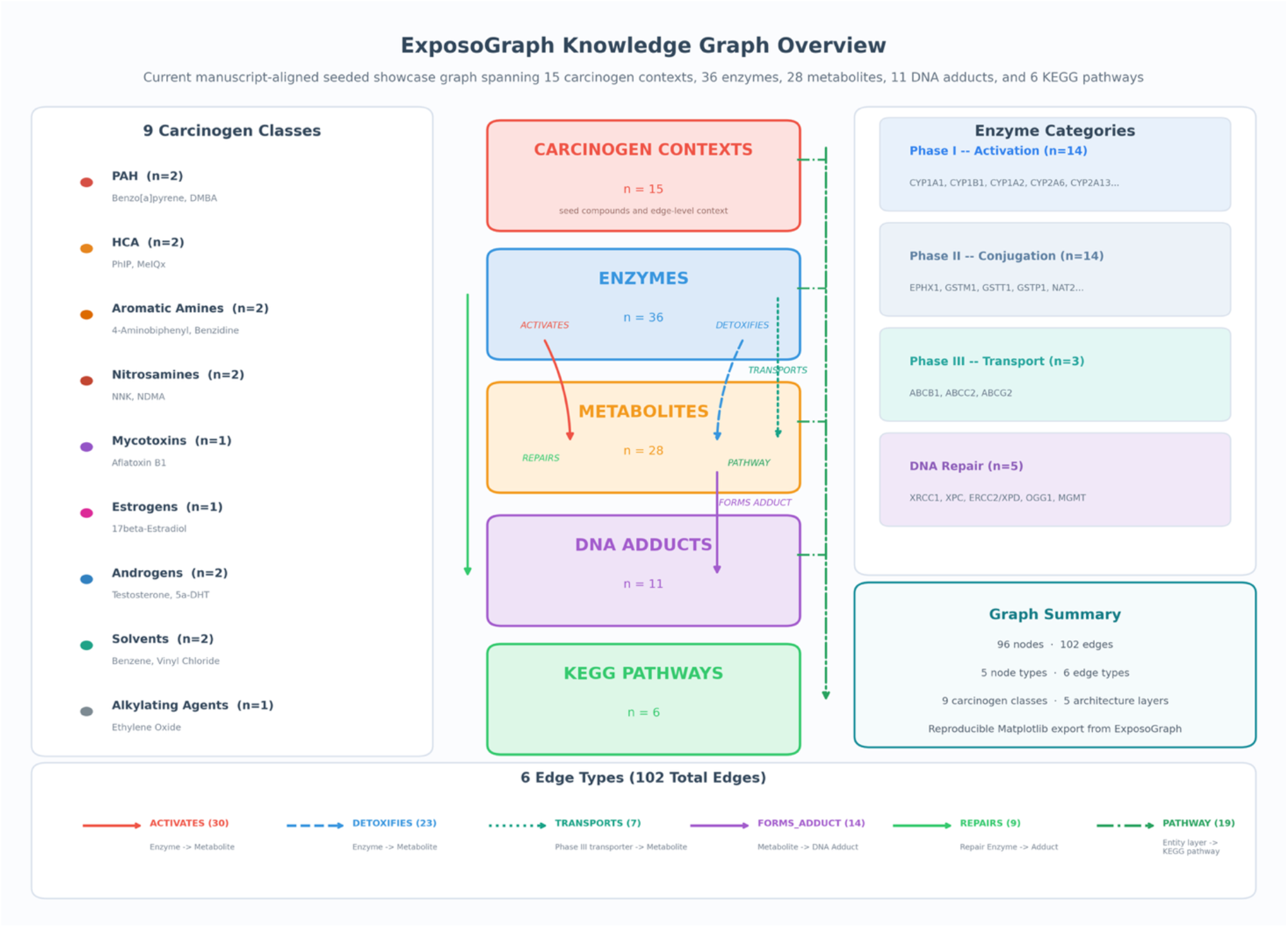
Overview of the current manuscript-aligned seeded ExposoGraph showcase graph. The figure summarizes curated carcinogen contexts across carcinogen classes, together with enzyme nodes spanning Phase I activation, Phase II conjugation, Phase III transport, and DNA repair. The overview also captures metabolite nodes, DNA adduct nodes, and KEGG pathway references, with edge families representing activation, detoxification, transport, adduct formation, repair, and pathway membership. All counts and grouped inventories are derived directly from the seeded showcase graph at export time.

### Visualization implementation

The ExposoGraph visualization layer is implemented through three complementary interfaces: rapid interactive previews generated with Plotly, standalone HTML exports rendered with a packaged D3-HTML viewer, and an advanced browser-based viewer implemented with Dash Cytoscape (22). Across these views, nodes are color-coded by semantic type: carcinogens in rose-red (#e05565), enzymes in teal-blue (#4f98a3), genes in dark teal (#3d8b8b), metabolites in amber-orange (#e8945a), DNA adducts in violet (#a86fdf), pathways in blue (#5591c7), and tissues, where present, in brown-gold (#c2855a). In the advanced viewer, node shape and size are also type-specific, with carcinogens rendered as diamonds, DNA adducts as hexagons, pathways as rounded rectangles, and tissues as tag-like nodes. Edge color is the primary encoding of biological function across views, and the Dash Cytoscape viewer adds further styling for selected relationship classes, including visually de-emphasized dashed pathway edges, dotted REPAIRS edges, thicker FORMS_ADDUCT edges, and directed arrows for supported relation types. The standalone D3 viewer uses a type-aware force-directed layout with differentiated link distances, repulsive forces, and collision handling, while the advanced viewer additionally supports force (COSE), hierarchical, circular, and saved-preset layouts.

The platform provides several layers of interactivity. Search supports rapid identification of graph entities, with the advanced viewer matching labels, identifiers, detail text, tissue, phenotype, PMID, and related metadata; matching results automatically expand to immediate neighbors while dimming unrelated regions of the graph. Carcinogen-class filtering isolates subgraphs relevant to specific exposure classes, and node-type and edge-type controls allow selective display of entity and relationship categories. Hover interactions reveal rich tooltips and local neighborhood context, while clicking a node or edge pins a detail panel showing annotations such as IARC group, phase, role, reactivity, tissue context, variant, phenotype, source database, evidence, and provenance. The advanced viewer also provides zoom, pan, reset-view controls, PNG and SVG export, and layout export as JSON for reproducible saved positioning. Plotly previews retain hover metadata, legend toggles, pan, and scroll-zoom, while the standalone D3 HTML viewer supports drag repositioning, search, carcinogen-class filters, node-type highlighting via legends, detail panels, and zoom controls in a responsive SVG workspace.

## RESULTS

### Knowledge graph overview

The completed ExposoGraph contains 96 nodes and 102 directed edges spanning 9 carcinogen classes (Figure 2). The graph provides coverage of 15 carcinogen context nodes, 36 metabolizing enzymes and repair proteins (14 Phase I, 14 Phase II, 3 Phase III, and 5 DNA repair), 28 metabolic intermediates, 11 DNA adduct types, and 6 pathway nodes. Enzymes in the Phase I category include members of the CYP1, CYP2, CYP3, and CYP17/19 families, along with steroid reductases (SRD5A1, SRD5A2), aldo-keto reductase (AKR1C3), and HSD3B2. Phase II enzymes span major conjugation and detoxification families, including glutathione S-transferases (GSTM1, GSTT1, GSTP1), N-acetyltransferases (NAT1, NAT2), UDP-glucuronosyltransferases (UGT1A1, UGT2B7, UGT2B17, UGT2B15), sulfotransferase (SULT1A1), and additional detoxification enzymes (EPHX1, NQO1, COMT, AKR1C2). The DNA repair module includes base excision repair enzymes (OGG1, XRCC1), nucleotide excision repair proteins (XPC, ERCC2/XPD), and direct reversal repair (MGMT).

**Figure 2.**
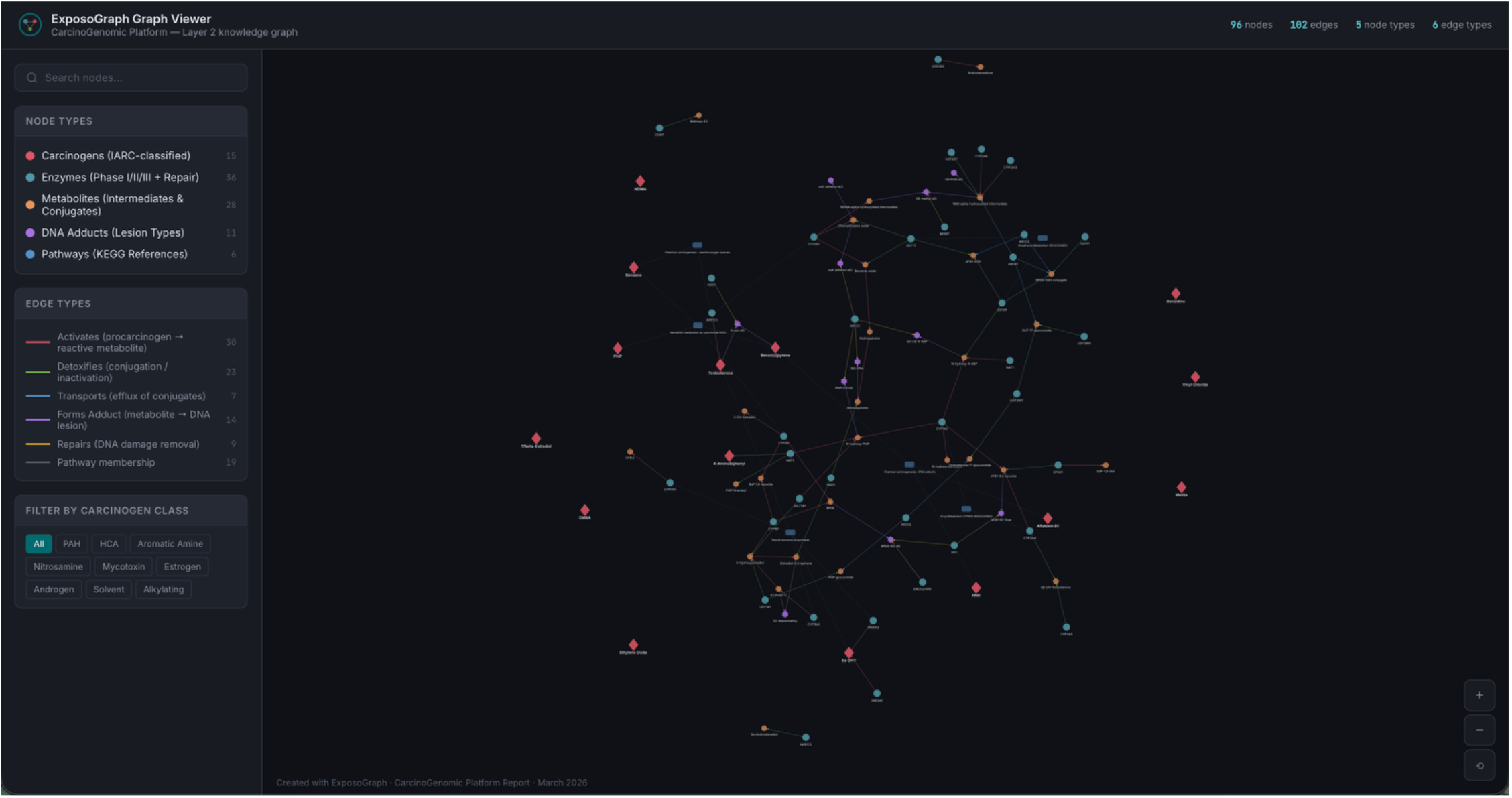
Example pathway visualization using ExposoGraph Viewer (D3). All carcinogen classes are visualized in this example graph.

### Carcinogen-specific pathway visualization

The ExposoGraph enables visualization of curated carcinogen-specific metabolic routes together with activation, detoxification, DNA adduct formation, and selected pharmacogenomic context. We illustrate this capability with four exemplar pathways (Figure 3A–D) rendered directly from the current ExposoGraph implementation, highlighting the diversity of activation mechanisms and the way genetic variation can be layered onto pathway interpretation.

**Figure 3.**
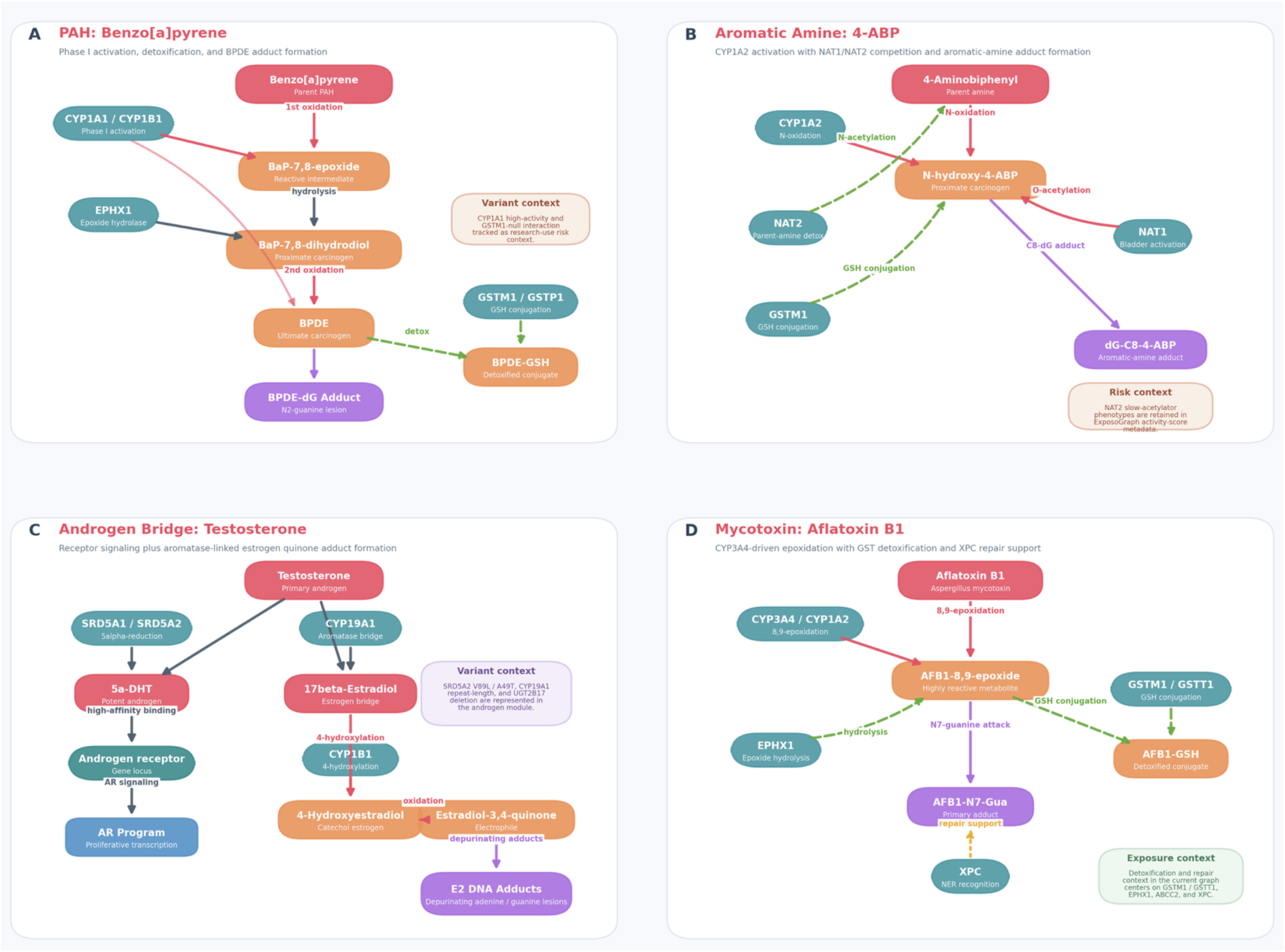
Exemple carcinogen-specific pathway panels rendered directly from ExposoGraph. The figure summarizes seeded BaP, 4-ABP, androgen-bridge, and aflatoxin B1 routes while keeping the panel content aligned to the current graph implementation.

The polycyclic aromatic hydrocarbon pathway (Figure 3A), exemplified by benzo[a]pyrene (BaP), involves initial activation by CYP1A1 and CYP1B1 to BaP-7,8-epoxide, followed by hydrolysis by EPHX1 to BaP-7,8-dihydrodiol and a second oxidation step yielding the ultimate carcinogen benzo[a]pyrene diol epoxide (BPDE). BPDE forms the BPDE-dG DNA adduct, while GSTM1 and GSTP1 support detoxification through glutathione conjugation to BPDE-GSH. In the current ExposoGraph figure, CYP1A1 high-activity and GSTM1-null states are shown as research-use variant context for this pathway (15).

The aromatic amine pathway (Figure 3B), represented by 4-aminobiphenyl (4-ABP), proceeds through CYP1A2-mediated N-oxidation to form N-hydroxy-4-ABP, a proximate carcinogenic intermediate. ExposoGraph highlights the competing roles of NAT2 in parent-amine detoxification and NAT1 in activation of the N-hydroxylated intermediate, together with GSTM1-linked detoxification context, culminating in the aromatic-amine DNA adduct dG-C8-4-ABP. NAT2 slow-acetylator phenotypes are retained in the ExposoGraph activity-score metadata as pathway-relevant pharmacogenomic context (13, 23).

The androgen metabolism module (Figure 3C) was incorporated to capture two complementary mechanisms by which androgens may contribute to carcinogenesis, particularly in the prostate. In one branch, testosterone is converted to 5-alpha-dihydrotestosterone (DHT) by SRD5A1 and SRD5A2, after which DHT engages the androgen receptor (AR) and downstream proliferative signaling. In the second branch, CYP19A1 converts testosterone to 17-beta-estradiol, which is then 4-hydroxylated by CYP1B1 and oxidized to estradiol-3,4-quinone, producing depurinating adenine and guanine DNA adducts. This aromatase bridge creates an explicit connection between androgen and estrogen-related carcinogenic metabolism in ExposoGraph, and the current module also annotates SRD5A2 V89L/A49T, CYP19A1 repeat-length variation, and UGT2B17 deletion as variant context (24, 25).

The mycotoxin pathway (Figure 3D), represented by aflatoxin B1 (AFB1), proceeds through CYP3A4- and CYP1A2-mediated 8,9-epoxidation to form the highly reactive AFB1-8,9-epoxide. This electrophilic intermediate can be detoxified through GSTM1/GSTT1-mediated glutathione conjugation and EPHX1-linked hydrolysis, or it can attack guanine to form the AFB1-N7-Gua adduct. In the current ExposoGraph implementation, repair support for this route is represented by XPC, and the panel emphasizes the balance between bioactivation, detoxification, and adduct handling rather than an extended downstream AFB1-FAPY repair cascade (26, 27).

### Interactive feature evaluation

The interactive ExposoGraph viewer supports multiple modes of exploration, including carcinogen-class filtering, node- and edge-type filtering, free-text search, hover tooltips, and click-to-inspect detail panels. Class-based filtering isolates the subgraph associated with a specific carcinogen group; for example, in the current showcase graph, selecting the Androgen filter retains 26 nodes and 19 edges while dimming unrelated graph content (Figure 4). Search is substring-based rather than autocomplete and matches labels, identifiers, detail text, group labels, tissue annotations, variants, phenotypes, and related metadata. Entering CYP1 highlights CYP1-family enzymes together with connected neighborhood context rather than restricting the view to exact family-name matches alone. Clicking or hovering a node highlights its local neighborhood and displays contextual annotations such as phase, role, detail text, tissue context, source database, and available provenance. For example, CYP1A1 is annotated in the current viewer as a Phase I activation enzyme with the detail PAH diol-epoxide formation; AhR-inducible; extrahepatic expression. Allele-level research-use activity summaries exist in the underlying ExposoGraph reference data, but they are not currently rendered in the viewer as rsID-specific tooltip effect sizes.

**Figure 4.**
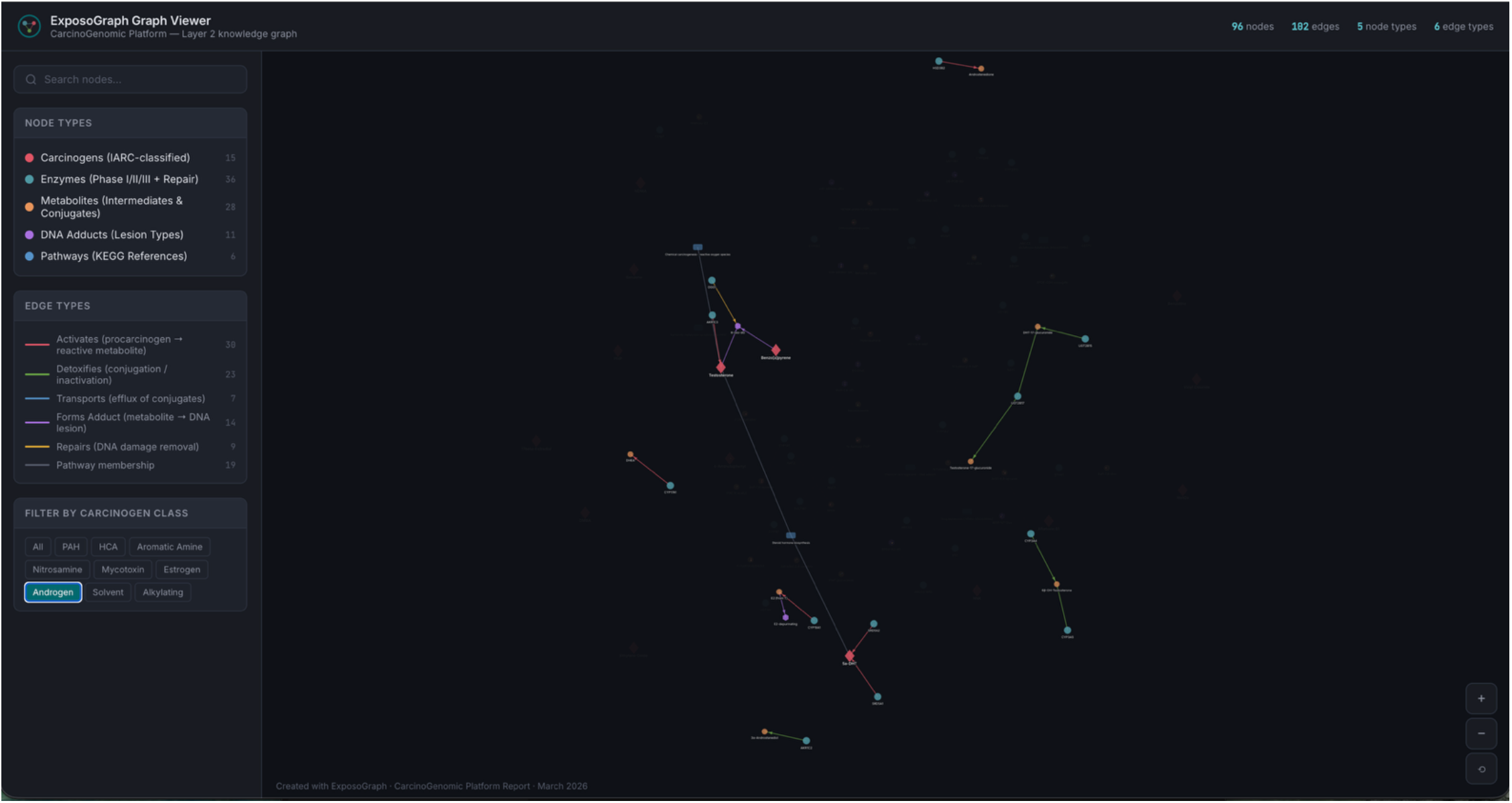
Example Androgen pathway visualization using ExposoGraph Viewer (D3). Only Androgen carcinogen class is visualized.

### Integration with pharmacogenomic risk scoring

The pathway structure encoded in ExposoGraph provides a foundation for downstream pharmacogenomic risk scoring. In the current implementation, a subset of enzyme nodes is annotated with representative activity-score values derived from curated allele-function tables, together with graph topology that distinguishes activation, detoxification, mixed-function, and repair roles. For guideline-backed pharmacogenes, activity-score conventions follow CPIC-aligned resources, whereas many carcinogen-metabolism genes currently use research-use literature-derived or database-backed summaries rather than fully standardized CPIC scoring. These annotations support class-level summaries of activation and detoxification capacity across carcinogen groups and enable topology-aware per-gene impact scoring based on each enzyme’s position relative to downstream DNA adduct formation and repair. In this way, ExposoGraph provides the structural framework for pharmacogenomic interpretation and future composite risk modeling, while the current Figure 5 is intentionally presented as a pharmacogenomic risk-scoring foundation rather than a full patient-level automated risk classifier.

**Figure 5.**
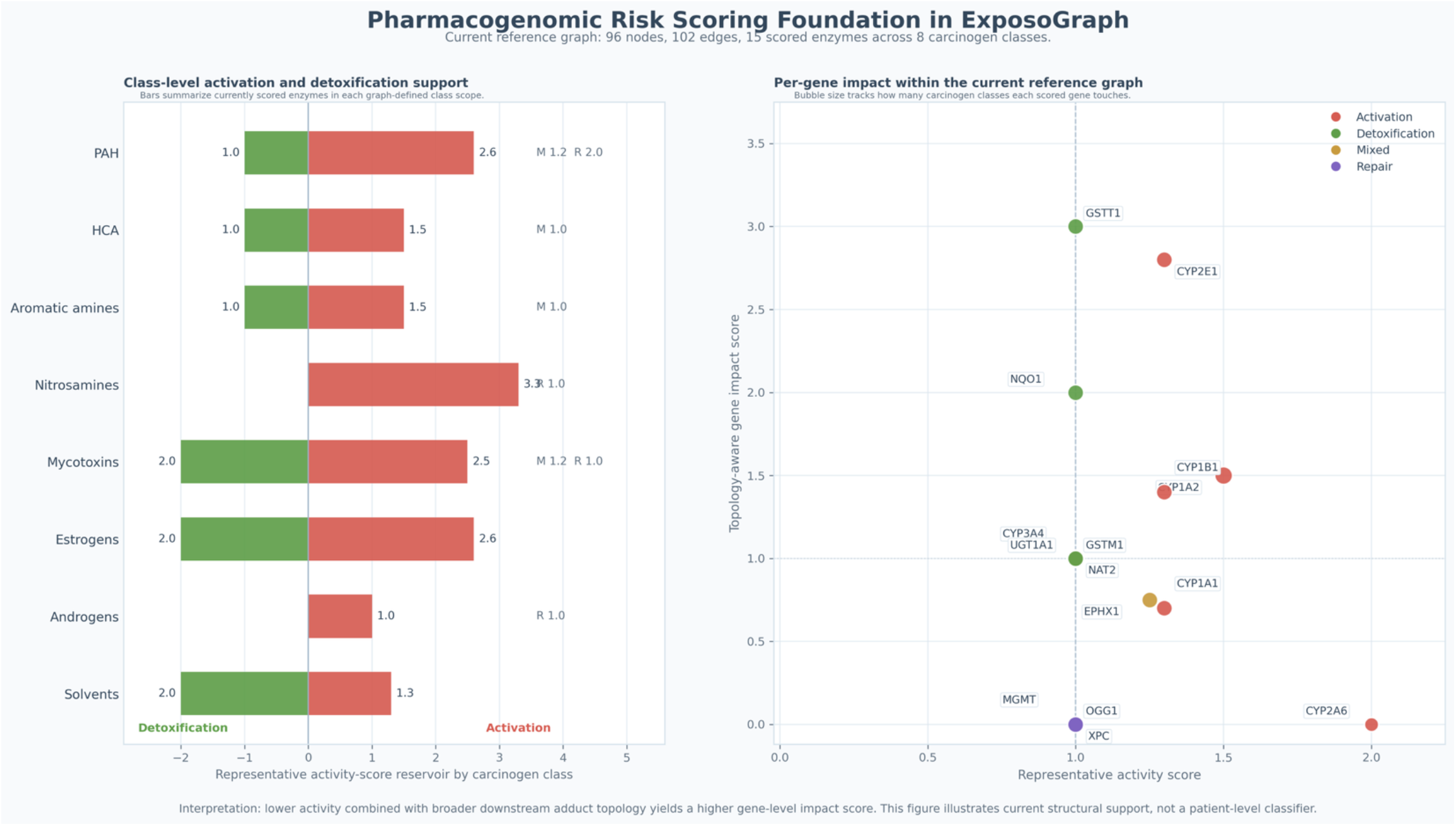
Pharmacogenomic risk-scoring foundation in ExposoGraph. In the current reference graph (96 nodes, 102 edges), 13 scored enzymes span 8 carcinogen classes. Left, horizontal bars summarize the current class-level reservoir of scored activation and detoxification support within each graph-defined carcinogen class. Positive red bars indicate summed representative activity scores for activation enzymes, whereas negative green bars indicate summed representative activity scores for detoxification enzymes; mixed-function (M) and repair (R) contributions are shown as text annotations where present. Right, each point represents a scored enzyme positioned by representative activity score on the x-axis and topology-aware gene impact score on the y-axis, where the impact score reflects downstream adduct-forming and repair reachability in the graph. Point color denotes role bucket (activation, detoxification, mixed, or repair), and bubble size reflects how many carcinogen classes each scored gene touches. This figure summarizes current structural support for pharmacogenomic interpretation in ExposoGraph and should be interpreted as a foundation for future composite risk modeling rather than a patient-level automated risk classifier.

### Comparison with existing tools

ExposoGraph addresses a gap not directly covered by existing bioinformatics platforms. The Comparative Toxicogenomics Database (CTD) provides a comprehensive catalog of curated chemical-gene-disease relationships and supports network-oriented exploration, but it is not organized as a pre-curated carcinogen-metabolism platform with integrated pharmacogenomic activity annotation and carcinogen-class filtering (18). KEGG provides manually curated reference pathway maps and interactive pathway-viewer tools, but it is not designed to support carcinogen-class exploration or genotype-informed pharmacogenomic interpretation (19). PharmCAT operates at the genotype-to-phenotype and prescribing-recommendation level for clinical pharmacogenomics but does not address carcinogen metabolism or cancer-focused pathway analysis (16, 17). Cytoscape provides a powerful general-purpose platform for network visualization and analysis with extensive app support, but building a carcinogen-specific knowledge graph in Cytoscape still requires substantial manual assembly and curation (20). ExposoGraph combines curated carcinogen-metabolism content, interactive browser-style visualization, and pharmacogenomic annotation in a single workflow, with topology-aware risk-scoring support that provides a foundation for future composite cancer-risk modeling.

## DISCUSSION

The CarcinoGenomic Knowledge Graph provides a framework for visualizing relationships among carcinogen exposure, genetic variation in metabolizing enzymes, and DNA damage. By integrating curated metabolic trajectories linking nine carcinogen classes, 36 enzymes, and 11 DNA adduct types within an interactive network, the ExposoGraph supports hypothesis generation, pathway exploration, and future risk-model development without requiring users to manually synthesize information across multiple resources.

The addition of androgen metabolism extends the framework beyond exogenous exposures and illustrates how endogenous hormone metabolism can intersect with genotoxic pathways. As shown in Figure 3, CYP19A1 (aromatase) links androgen and estrogen metabolism, providing a mechanistic route by which testosterone may contribute to carcinogenesis through both receptor-mediated proliferation after conversion to dihydrotestosterone (DHT) and genotoxic DNA damage after aromatization to estradiol and subsequent catechol estrogen quinone formation (24, 25). This dual-pathway view may be relevant to prostate cancer biology: variation in SRD5A2 may influence the proliferative branch, variation in CYP19A1 may influence the estrogen-linked genotoxic branch, and variation in UGT2B17 may affect overall androgen clearance. By representing these connected processes within a single network, the ExposoGraph may help researchers evaluate how combinations of variants across the androgen pathway contribute to testable prostate cancer risk hypotheses.

The ExposoGraph has several potential research and educational applications. First, it can generate testable hypotheses for gene-environment interaction (GxE) studies by highlighting enzyme-carcinogen pairs for which functional genetic variation is known but epidemiologic interaction data remain limited. Recent advances in cancer GxE analysis, including studies of colorectal cancer risk across multiple environmental exposures, underscore the importance of more systematic GxE characterization (28). Second, when paired with genotyping or whole-genome sequencing data, the platform could support exploratory individualized risk modeling by providing a visual and quantitative framework for interpreting how an individual’s metabolic enzyme profile may modify susceptibility to specific exposures. Third, the interactive network could serve as a useful teaching resource for chemical carcinogenesis by helping students and trainees trace metabolic pathways and understand the mechanistic basis of GxE relationships. Fourth, the platform may also help prioritize questions for occupational and environmental health research by visualizing how genetic profiles and workplace exposures may intersect.

The ExposoGraph is also positioned to interface with emerging computational and genomic resources. Pathway-specific polygenic risk scores (pPRS) that weight variants according to their functional roles in carcinogen metabolism could leverage the graph topology to define biologically coherent variant sets and pathway boundaries (29). Integration with large-scale genomic resources such as the All of Us Research Program could enable population-level evaluation of hypotheses generated from the graph (30). GTEx-derived tissue-expression annotations provide a complementary resource for identifying tissues in which metabolizing enzymes are most highly expressed, thereby informing tissue-specific risk models (21). Population allele-frequency data from gnomAD can further contextualize how common potentially relevant variants are across ancestral populations (31, 32). Established pharmacogenomic frameworks such as those reviewed by Daly provide a translational precedent for how curated gene-variant-function knowledge may eventually be incorporated into clinical workflows (3).

Several limitations should be acknowledged. First, the current graph represents a curated subset of carcinogens with relatively well-characterized metabolic activation pathways; expansion to include additional IARC Group 1 and Group 2A agents remains ongoing. Second, the enzyme activity scores used in composite risk classification are derived from literature-based estimates from in vitro and epidemiologic studies and would benefit from standardization through large-scale functional assays. Third, the current visualization does not model dynamic modifiers of metabolic capacity, such as enzyme induction or inhibition by co-exposures, which can substantially alter pathway flux. Fourth, population-level validation of any composite risk framework will require prospective cohorts with both genotyping data and quantified exposure assessment, which remain logistically challenging to assemble.

Future development will expand the ExposoGraph to include additional carcinogen classes and mechanistic modules, including heavy metals, ionizing radiation, and biological agents such as oncogenic viruses. We also plan to deepen the incorporation of tissue-specific gene-expression data to model organ-specific activation patterns more explicitly and to explore machine-learning approaches, including graph neural networks trained on ExposoGraph topology, to assess whether the network can help predict novel carcinogen-enzyme relationships and prioritize new GxE candidates for experimental and epidemiologic follow-up.

As precision medicine advances, understanding how inherited variation interacts with environmental exposure profiles will become increasingly important for cancer prevention. The CarcinoGenomic Knowledge Graph provides an interactive platform for visualizing these relationships by integrating curated information from IARC, KEGG, PharmVar, CPIC, CTD, and GTEx-derived annotations into a unified network.

By bringing these resources together, the platform addresses an important gap in carcinogenomic research. Its interactive design, pharmacogenomic annotation, and support for downstream modeling provide a foundation for future work on risk stratification, hypothesis generation, and education focused on the interplay between genetic susceptibility and carcinogen exposure. We anticipate that it will be useful to researchers, clinicians, educators, and public health professionals working to better understand environmentally mediated cancer.

## Acknowledgement

Assistance with language editing, grammatical revision, and correction of typographical errors was provided through the use of artificial intelligence tools, including ChatGPT 5.4, Claude Opus 4.6, Microsoft 365 Copilot, and Perplexity. The authors are responsible for all content. All authors have seen and approved the manuscript, and that it hasn’t been accepted or published elsewhere.

## CRediT authorship contribution statement

K.J.P: Conceptualization, Writing – original draft, review & editing, Funding acquisition. J.U.K.: Conceptualization, Writing – original draft, review & editing.

## Code availability

The source code is available through GitHub (https://github.com/kazilab/ExposoGraph), with documentation provided at https://exposograph.readthedocs.io. The package is also available through PyPI (https://pypi.org/project/ExposoGraph/) and web browser at Streamlit (https://exposograph.streamlit.app).

